# An adaptor for feedback regulation of heme biosynthesis by the mitochondrial protease CLPXP

**DOI:** 10.1101/2024.07.05.602318

**Authors:** Thomas Cottle, Lydia Joh, Cori Posner, Adam DeCosta, Julia R. Kardon

## Abstract

Heme biosynthesis is tightly coordinated such that essential heme functions including oxygen transport, respiration, and catalysis are fully supplied without overproducing toxic heme precursors and depleting cellular iron. The initial heme biosynthetic enzyme, ALA synthase (ALAS), exhibits heme-induced degradation that is dependent on the mitochondrial AAA+ protease complex CLPXP, but the mechanism for this negative feedback regulation had not been elucidated. By biochemical reconstitution, we have discovered that POLDIP2 serves as a heme-sensing adaptor protein to deliver ALAS for degradation. Similarly, loss of POLDIP2 strongly impairs ALAS turnover in cells. POLDIP2 directly recognizes heme-bound ALAS to drive assembly of the degradation complex. The C-terminal element of ALAS, truncation of which leads to a form of porphyria (XLDPP), is dispensable for interaction with POLDIP2 but necessary for degradation. Our findings establish the molecular basis for heme-induced degradation of ALAS by CLPXP, establish POLDIP2 as a substrate adaptor for CLPXP, and provide mechanistic insight into two forms of erythropoietic protoporphyria linked to CLPX and ALAS.

## MAIN TEXT

Mitochondria sustain and control the activities of their proteins using AAA+ unfoldases. The mitochondrial AAA+ unfoldase CLPX, which forms a proteolytic complex with the protease CLPP (CLPXP), is essential to mitochondrial function. Perturbation of CLPXP function underlies several mitochondriopathies and drives cancer progression (*1–3*). Unlike the related 26S proteasome, CLPX and other mitochondrial unfoldases recognize substrates without ubiquitination. These unfoldases thus require alternative and largely undescribed mechanisms for specific and conditional recognition of their substrates.

In one important role, CLPX exerts bidirectional control over an essential biosynthetic function of mitochondria, heme biosynthesis, by acting on the first and rate-limiting enzyme, ALA synthase (ALAS). In addition to performing oxygen transport as part of hemoglobin in vertebrate red blood cells, heme is required as an enzymatic cofactor and sensor molecule. Underproduction of heme impairs these functions, whereas overstimulation of heme biosynthesis cause accumulation of toxic heme precursor and iron depletion (*4*) Heme biosynthesis is thus heavily regulated, particularly at its entry point, ALAS; impaired function or regulation of ALAS causes several forms of anemia and porphyria (*5*). We previously discovered that CLPX activates ALAS through partial unfolding to facilitate cofactor incorporation, supporting heme biosynthesis and erythropoiesis (*6*, *7*). Heme-replete conditions in cells also induce CLPX-dependent degradation of ALAS for both housekeeping (ALAS1) and erythroid (ALAS2) isoforms (*3*, *8*), suggesting that heme feedback redirects CLPXP to complete unfolding and degradation of ALAS. This degradation requires a putative heme-binding motif (CP) located within the flexible N-terminus that extends from the folded enzyme core of both ALAS isoforms (*8*). Degradation was not reconstituted, however, by simple addition of heme (Fig. 1A), leaving open the question of how heme induces CLPXP to degrade ALAS and thus regulates its own synthesis. We hypothesized that CLPX might require a direct substrate adaptor to degrade ALAS. Several substrate adaptors have been identified for bacterial homologs of CLPX (*9*), but these proteins have no known homologs in eukaryotes.

**Figure 1:**
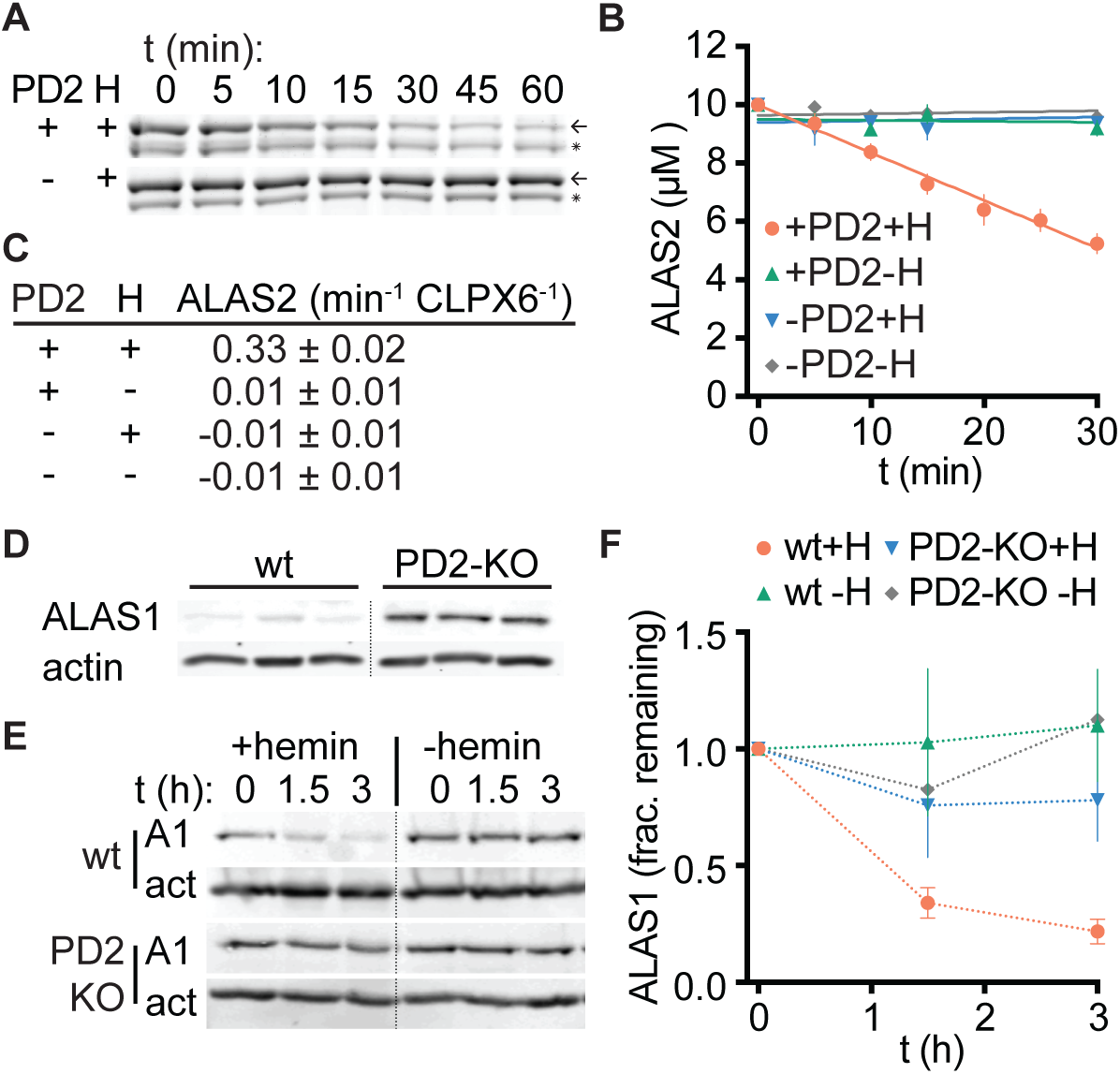
POLDIP2 is crucial for heme-induced degradation of ALAS by CLPXP. (**A**) Representative images of ALAS2 degradation time courses *in vitro*. 10 µM ALAS2 was incubated with 0.5 µM CLPX and CLPP (hexamer and 14-mer respectively), 4 mM ATP, 3 µM POLDIP2 (PD2), +/- 10 µM hemin (H). (n = 5 for +hemin + PD2, 3 for other conditions). Samples were separated by SDS-PAGE and visualized by Sypro Red stain. Upper band (highlighted with arrow) is full-length ALAS2; lower band (asterisk) corresponds to C-terminally proteolyzed ALAS2 fragment (*14*). See Fig. S1 for additional gel images +/-POLDIP2. (**B**) Plot of degradation of ALAS2 *in vitro* as monitored in (A). Lines represent linear fits to data, error = SEM. (**C**) Rates extracted from (B. (**D**) Western blot of steady-state ALAS1 levels in wt or PD2-KO HEK293 cells. Samples from three independent cultures of each cell line are shown. (**E**) Degradation of ALAS1 in wt or PD2-KO HEK293 cells. Endogenous heme production was blocked with 0.5 mM succinylacetone 24 hours prior to observation. At t_0,_ translation was inhibited with 100 µg/mL cycloheximide and 100 µM hemin or vehicle (DMSO) was added. ALAS1 (A1) and actin (act) were detected by western blot. (**F**) Quantitation of ALAS1 levels from (E) (n = 3, error = SEM).

### POLDIP2 is an adaptor for heme-induced degradation of ALAS by CLPXP

DNA polymerase delta interacting protein 2 (POLDIP2) was initially characterized as an accessory factor for several error-prone nuclear DNA polymerases. It primarily localizes to the mitochondrial matrix, however, and critical contacts for activity with DNA polymerase are removed by mitochondrial targeting sequence cleavage, suggesting that it has a distinct function in the mitochondrion (*10*). POLDIP2 has also been identified as an interactor and possible regulator of CLPX (*11*, *12*), but the mechanism by which POLDIP2 might regulate CLPX activity has not been determined. Interaction between POLDIP2 and ALAS has also been detected by mass spectrometry, suggesting that POLDIP2 could participate in regulation of ALAS by CLPX (*8*, *13*). We therefore tested whether addition of POLDIP2 to the CLPXP degradation machinery could reconstitute degradation of ALAS by CLPXP. As we previously observed, purified human ALAS2 (the erythroid isozyme of ALAS) could not be degraded by CLPXP alone (*7*), even with the addition of heme. Addition of POLDIP2 in the presence of heme, however, induced robust degradation of ALAS2 (Fig. 1A-C; Fig. S1). The addition of POLDIP2 had no effect on degradation when heme was not present. Therefore, POLDIP2 is necessary and sufficient to reconstitute heme-responsive degradation of ALAS2 by CLPXP. We additionally noticed that a truncation product of ALAS2 (C-terminal based on our purification strategy and previous observations of bacterially expressed ALAS2 in other groups (*14*, *15*)) appeared to be much more slowly degraded, which we discuss further below.

To test the physiological requirement for POLDIP2 in the regulation of ALAS by heme, we monitored ALAS1 protein levels in wildtype and POLDIP2-knockout (PD2-KO) HEK293 cells. At steady state, ALAS1 levels in PD2-KO cells were dramatically increased relative to wildtype cells (Fig. 1D, Fig. S2), suggesting that endogenous heme levels induce turnover of ALAS1 that requires POLDIP2. We monitored the heme- and POLDIP2-dependence of ALAS1 degradation in cells by translational shutoff, depleting endogenous heme and equalizing ALAS1 protein levels by inhibiting heme biosynthesis with succinylacetone for 24 hours prior to observation. In both cell lines, little degradation is observed without heme addition. In wildtype cells, ALAS1 was degraded quickly after heme addition, nearing completion at 3 hours. In the POLDIP2 knockout line, degradation is dramatically reduced (Figure 1E,F). Residual degradation may be due to LONP, which has been previously implicated in ALAS1 degradation (*8*, *16*). These data confirm that POLDIP2 is critical for heme-responsive degradation of ALAS in cells. With our *in vitro* observation of POLDIP2-mediated ALAS2 degradation, these data also demonstrate that POLDIP2 is critically important for heme-feedback regulation of both the erythroid and housekeeping isozymes of ALAS.

One mechanism by which POLDIP2 might function is as a physical adaptor, tethering ALAS to CLPX. Supporting this idea, POLDIP2 directly interacted with ALAS2 in the presence of heme, as judged by co-immunoprecipitation with purified proteins. However, in the absence of heme no interaction was observed (Fig. 2A). Furthermore, POLDIP2 recruits ALAS2 to ATPγS-locked CLPXP. Whereas ALAS2 coprecipitated weakly with CLPXP alone, with some increase in the presence of heme, POLDIP2 strengthened CLPX-ALAS2 association modestly in the absence of heme and dramatically in its presence (Fig. 2B). As previously reported, POLDIP2 bound constitutively to CLPX (*12*); we observed no change in interaction due to heme addition. Together with our reconstitution of degradation, these data demonstrate that POLDIP2 functions as a heme-sensitive adaptor for ALAS, functioning at least in part to tether ALAS to CLPX. Although CLPX has some affinity for ALAS, as has been previously observed for yeast and human housekeeping orthologs (*7*, *17*), our data indicate that POLDIP2 is the primary site of heme-sensitive interaction with ALAS.

**Figure 2:**
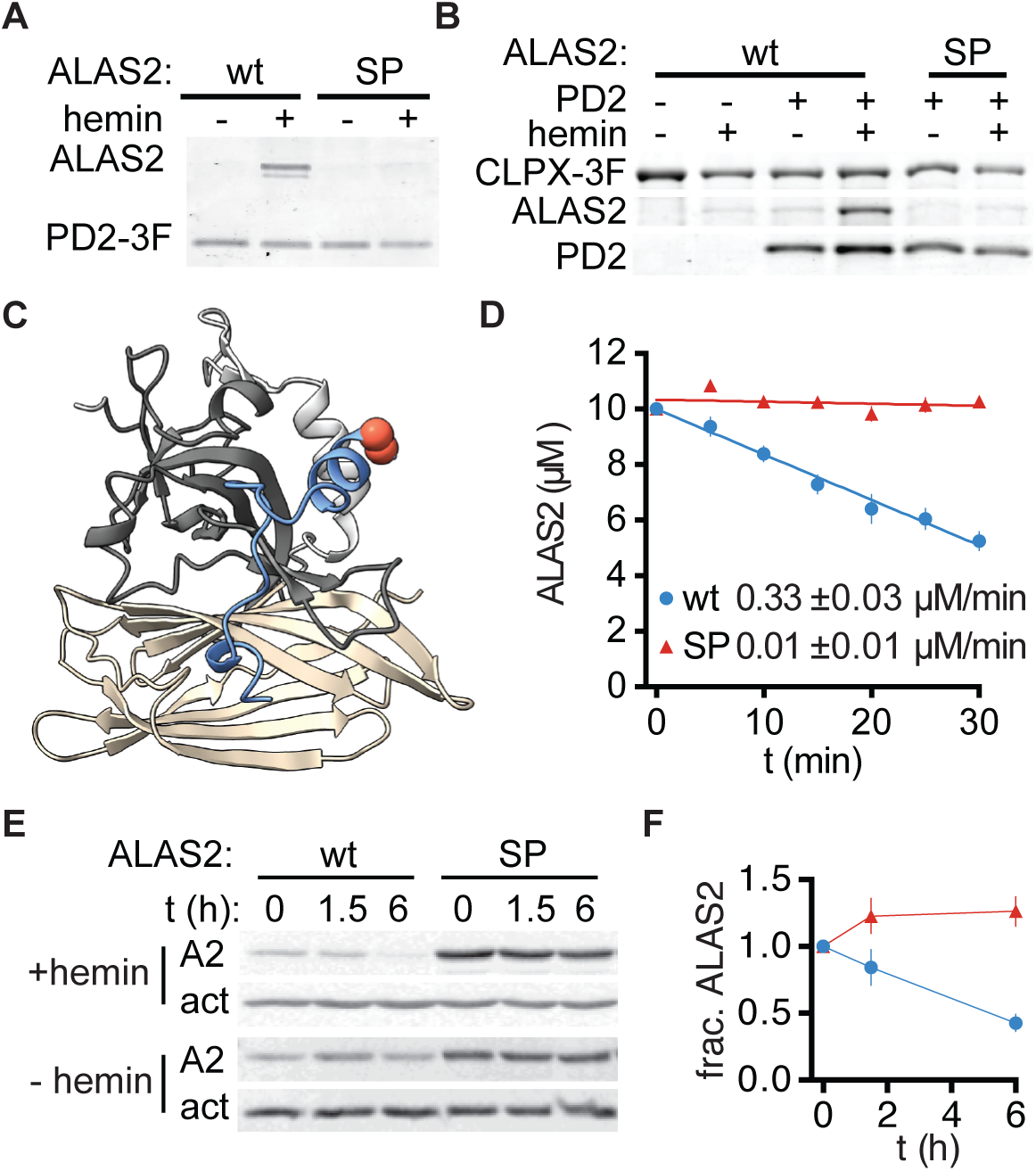
A heme-regulatory motif in ALAS drives direct ALAS2-POLDIP2 interaction and ALAS degradation. (**A**) Co-immunoprecipitation of 5 µM ALAS2 (wt or C70S, ‘SP’) with 5 µM POLDIP2-3xFLAG (PD2-3F), +/- 10 µM hemin. Samples were visualized in Sypro Red-stained SDS-PAGE gels. Representative image shown, n = 3. (**B**) Co-IPs performed as in (A), but with 0.5 µM hexamer ClpX-3xFLAG (CLPX-3F), 0.5 µM 14-mer CLPP, and 4 mM ATPγS. Representative image shown, n = 3. (**C**) Alphafold2 model of a POLDIP2-ALAS2 complex. POLDIP2 is shown in dark gray (YccV domain), light gray (linker), and wheat (ApaG domain). The interacting peptide of ALAS2 (C70-D95) is depicted in blue, with C70 and P71 highlighted with red spheres at the α carbon. (**D**) Plot and rates of *in vitro* degradation of ALAS2 and ALAS2^SP^ as in Figure 1, with POLDIP2 and hemin (n = 5 for wt, 3 for SP variant). (**E**) Degradation of transiently transfected 1D4-tagged ALAS2 variants in HEK293 cells. Cycloheximide and hemin or vehicle were added as in Figure 1C. ALAS2 and actin levels were detected by Western blot. (**F**) Quantitation of ALAS levels over time after addition of hemin in (E) (n = 3).

We used AlphaFold2-multimer with ColabFold (*18*) to generate a model for the ALAS2-POLDIP2 complex. POLDIP2 was predicted with high confidence (PAE <5 at core contact points) to interact with a ∼26-residue region in the disordered N-terminal element (*14*) of ALAS2 (Fig. 2C). This region includes the conserved CP motif (C70-P71) that is predicted to bind heme and is required for heme-responsive degradation of ALAS1 in cells (*8*). POLDIP2 is composed of a YccV and an ApaG domain, linked by an α-helical element. The POLDIP2-contacting element of ALAS2 contacts both domains of POLDIP2 and their linker, winding across one face of POLDIP2 in an extended conformation with some α-helical structure. No other contact between these proteins were predicted and due to the flexibility of the remainder of the N-terminal element of ALAS2, no significant orientation of the core of ALAS2 relative to POLDIP2 was observed. The involvement of the putative heme-binding motif of ALAS2 in the predicted interaction with POLDIP2 provides a structural rationale for why POLDIP2-ALAS2 binding is heme sensitive. POLDIP2 and CLPX were previously demonstrated to interact by their YccV and zinc-finger domains, respectively (*8*). AlphaFold2-multimer predicted that a short extension N-terminal to the YccV domain of POLDIP2 pairs with a Β sheet in the CLPX zinc finger. In a model of ALAS2, POLDIP2, and CLPX interaction generated by AlphaFold2, these contacts are recapitulated in a spatially compatible ternary complex (Fig. S3).

Instances of both domains that constitute POLDIP2, YccV and ApaG, recur in proteins controlling substrate selection among several evolutionarily distant AAA+ protease systems. The original YccV protein is a modulator of substrate selection for the proteobacterial AAA+ proteases Lon and ClpCP (*19*, *20*). In humans, the F-box family of E3 ligases includes both a YccV- and an ApaG-domain-containing member (FBXO21 and FBXO3), controlling substrate selection by the extraordinarily complex AAA+ protease, the 26S proteasome (*21*, *22*). The YccV-domain protein ClpF, an adaptor for the plastid ClpC1 protease complex, regulates heme-induced degradation of GluTR, the initiating enzyme in the C5 heme biosynthetic pathway used in plants, most bacteria, and archaea (*23*), thus fulfilling a strikingly similar function to POLDIP2, although GluTR is not homologous to ALAS2 and an anti-adaptor, rather than ClpF, appears to be directly regulated by heme (*24*). In an additional similarity to a bacterial adapter protein, the YccV domain is part of a subfamily of SH3-like domains that includes SspB, an adaptor protein for bacterial ClpX (H-group co-membership in ECOD (*25*)). POLDIP2, like SspB, also appears to bind ClpX through pairing of a strand extending from the SH3 domain (N-terminal for POLDIP2 (Fig. S3), C-terminal for SspB (*26*)) with the Β sheet of the ClpX zinc-finger domain. Future characterization will reveal how the structural similarities in each of these domains may relate to shared mechanisms for regulation of proteolysis.

### A heme-binding motif on ALAS drives assembly of the degradation complex

Based on the presence of the CP motif at the predicted ALAS2-POLDIP2 interface (*8*), we hypothesized that this motif would be critical for POLDIP2 to induce degradation of ALAS2 by CLPXP in the presence of heme. ALAS2 with a mutation in this motif (C70S, ALAS2^SP^) did not coprecipitate with FLAG-tagged POLDIP2 in the presence or absence of heme and its interaction with CLPXP and POLDIP2 was not stabilized by heme (Fig. 2A, B). ALAS2^SP^ was not degraded in our reconstituted CLPXP-POLDIP2 system (Fig. 2D). To confirm the importance of the CP motif for degradation of ALAS2 in cells, we transiently transfected epitope-tagged ALAS2 variants into HEK293 cells and monitored degradation after translational arrest. Addition of heme induced robust degradation of wildtype ALAS2, but ALAS2^SP^ was not degraded (Fig. 2E, F). The CP motif of ALAS is thus essential for POLDIP2 to orchestrate assembly of a complex for heme-induced degradation.

### ALAS2 and POLDIP2 form a heme-binding complex

Based on our finding that heme drives POLDIP2 to bind ALAS and commit it for degradation, we sought to determine how heme is sensed within the ALAS-POLDIP2 complex. Due to the importance of the N-terminal CP motif in ALAS for heme-induced interaction with POLDIP2 and degradation, we expected that this motif would bind heme by direct coordination of iron. The porphyrin ring of heme supplies four ligands to its bound iron atom; heme-binding proteins can provide one or two axial ligands, inducing distinct shifts in the Soret band of heme. In the presence of ALAS2, the absorbance peak of heme shifted from near 390 nm to 375 nm, indicating direct binding (Fig. 3B). This absorbance shift is strongly correlated with a single iron ligand provided by a CP motif (*27*).

**Figure 3:**
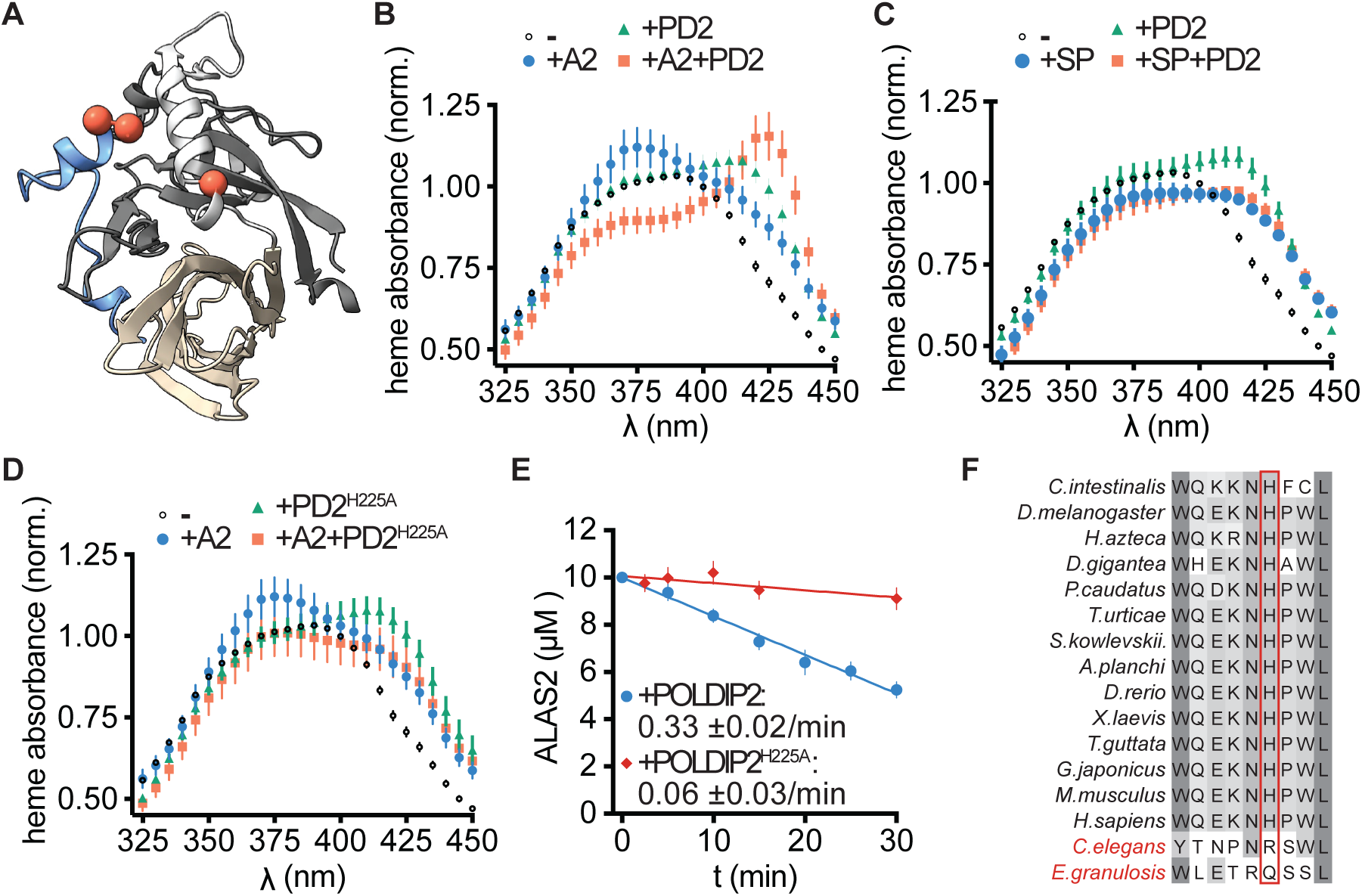
POLDIP2 and ALAS2 form a heme-coordinating complex. (**A**) Alphafold2 model of the POLDIP2-N-ALAS2 as in Figure 2A, where POLDIP2 is grey, the N-terminus of ALAS is blue, with ALAS C70-P71 and POLDIP2 H225 shown as red spheres at the α carbon. (**B**) UV-vis absorbance spectra of heme +/-ALAS2 (A2), POLDIP2 (PD2), or both (25 µM each protein and heme). Protein-only absorbance was subtracted and data were normalized to heme-alone absorbance at 390 nm. (**C**) Absorbance spectra as in (B), with ALAS2^SP^ (SP) instead of wildtype ALAS2. (**D**) Absorbance spectra as in (B), with POLDIP2^H225A^ (H225A) instead of wildtype POLDIP2. Heme-alone reference spectra are the same as in (C). (**E**) Degradation of ALAS2 *in vitro* as in Fig. 1B, with either POLDIP2 or POLDIP2^H225A^ and hemin (n = 5 for wt, 3 for H225A variant). (**F)** Sequence alignment of POLDIP2 near modeled heme-interacting site, with H225 position boxed in red. Heme-auxotrophic nematode species names are in red. N = 3 and error = SEM for absorbance and degradation data.

Because POLDIP2 is predicted to bind near the CP motif of ALAS (Fig. 3A), we hypothesized that POLDIP2 could directly co-coordinate the heme iron with ALAS. POLDIP2 induced a red shift in heme absorbance, indicating interaction between POLDIP2 and heme. In the presence of both POLDIP2 and ALAS2, heme absorbance shifted to a strong peak near 425 nm (Fig. 3B). This UV-vis absorbance profile is strongly correlated with heme iron coordinated by both a CP motif and another protein ligand (*27*), suggesting that ALAS2 and POLDIP2 co-ligate heme at the complex interface. ALAS2^SP^ failed to induce either a blue shift in heme absorbance on its own or a 425-nm peak with POLDIP2, indicating the N-terminal CP motif as the primary site of heme binding to ALAS2 (Fig. 3C).

We sought to identify a residue in POLDIP2 that could co-coordinate heme with ALAS2. Inspection of our model of the POLDIP2-ALAS2 complex revealed a histidine (H225), a common heme-iron-coordinating residue (*27*), in the linker between the POLDIP2 YccV and ApaG domains, near the ALAS2 CP motif (Fig. 3A). We mutated this residue to alanine (POLDIP2^H225A^) and tested its interaction with heme by UV-vis spectroscopy. POLDIP2^H225A^ in the presence of ALAS2 failed to induce a 425 nm peak in the heme absorbance profile, indicating that this residue is necessary for the formation of the ALAS2-POLDIP2 heme coordination complex (Fig. 3D). POLDIP2^H225A^ abolished the 375 nm peak in heme absorbance observed with ALAS2 alone, instead exhibiting absorbance similar to ALAS2^SP^, which does not bind heme. This suggests that POLDIP2^H225A^ can form a complex with ALAS2 that excludes heme from binding ALAS2, perhaps driven by the high concentrations of protein used in this equilibrium method (25 µM, compared to 5-10 µΜ in the nonequilibrium coIP experiments). On its own,

POLDIP2^H225A^ induced Soret band shifts equivalent to those of wild-type POLDIP2, suggesting that this intrinsic heme interaction is distinct from that involved in recognition of heme-bound ALAS. Supporting the importance of this position for degradation, we found that POLDIP2^H225A^ substantially impairs *in vitro* degradation of ALAS2 by CLPXP (< 20% of wild type rate (Fig. 3E), although binding to CLPX was unimpaired (Fig. S4). Supporting the importance of the POLDIP2 H225 region to recognition of heme-ALAS2, this residue and those immediately proximal to it are highly conserved among POLDIP2 homologs, with the notable exception of heme-auxotrophic nematodes (Fig. 3F). These organisms have lost heme biosynthetic genes, including ALAS homologs, and therefore would not require residues in POLDIP2 specific to recognition of ALS. These data together demonstrate that ALAS2 and POLDIP2 together form a distinctive coordination complex with heme and that this coordination is required for complex formation and subsequent degradation of ALAS2 by CLPXP and POLDIP2.

### N- and C-terminal elements of ALAS2 differentially direct degradation complex assembly and licensing

The reduced degradation we observed for the C-terminal ALAS2 truncation product (Fig. 1A) led us to consider whether the C-terminal element of ALAS2 is important for degradation. The C-termini of eukaryotic ALAS homologs are extended beyond the core enzyme fold shared with bacterial homologs. In crystal structures of human ALAS2, the ∼40 amino acid C-terminal extension is in an extended conformation along the exterior of the same protomer, positioning a small alpha helical segment to partially block the active site (Fig. 4A) (*14*). To test whether this element is important for degradation of ALAS2 by CLPXP, we generated C-terminally truncated variants of ALAS2 (Fig. 4B) and tested their susceptibility to degradation by CLPXP. Degradation was nearly ablated by truncation of the entire C-terminal region of ALAS (ALAS2^ΔC^) and was substantially slowed by truncation of the flexible C-terminal twelve amino acids (ALAS^Δ576^, depicted as yellow element in Fig. 4A,B) (Fig. 4C). Binding of ALAS2^ΔC^ to POLDIP2 was similar to that of wildtype, indicating that the C-terminus of ALAS2 is important for degradation but not for complex assembly (Fig. 4D). Because unfolding by CLPX is often initiated at a terminus (*28*), we compared the effect of occlusion of the ALAS2 termini. Occlusion of the C-terminus by fusion with a folded domain (SUMO) impaired degradation, whereas N-terminally occluded ALAS2 was readily degraded by the POLDIP2-CLPXP complex in the presence of heme (Fig. 4B,E; Fig. S5). These data demonstrate that the N-terminal element of ALAS2 is required for complex assembly but does not need to be free to induce degradation, whereas the C-terminal element of ALAS2 is dispensable for complex assembly but required for degradation. Several alleles of ALAS2 that truncate the C-terminal extension lead to a form of erythropoietic protoporphyria (XLDPP) (*29*, *30*). Although this disorder is likely due in part to the several-fold increase in enzyme activity attributed to loss of autoinhibition by the C-terminus (*14*, *15*, *30*, *31*), a C-terminally truncated ALAS2 variant also exhibited reduced turnover in cells (*32*). We propose that loss of degradation by CLPXP exacerbates the excess of ALAS2 activity in XLDPP. (*14*, *15*, *29*, *31*)

**Figure 4:**
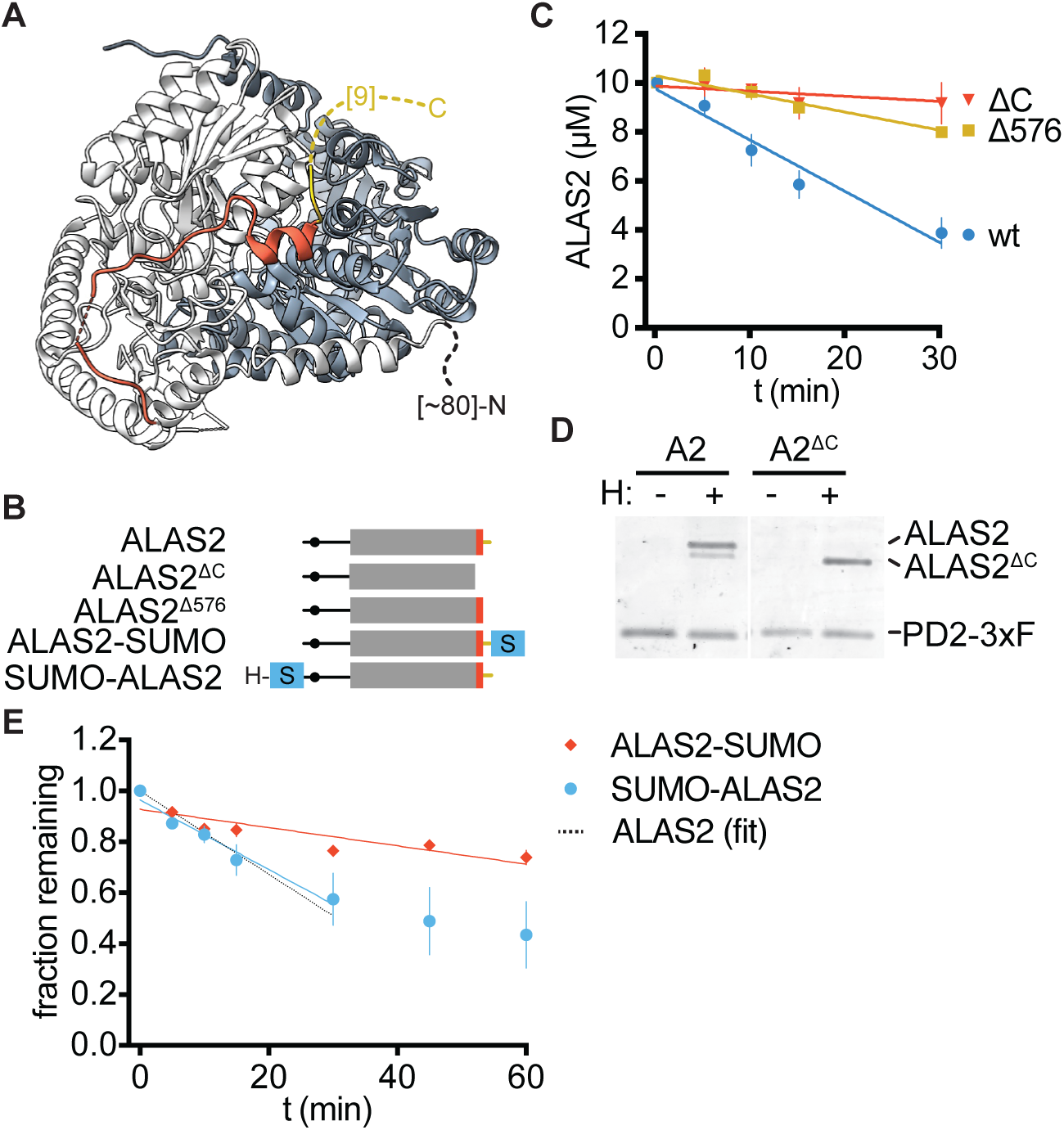
The N- and C-terminal elements in ALAS form functionally distinct contacts during heme-induced degradation by CLPX. (**A**) ALAS2 structure (PDB: 6HRH) with the C-terminal extension highlighted on one protomer in orange (docked portion of the C-terminus, G544-F575) and yellow (project end, G576-M578). The nine C-terminal residues (G579-A587) are disordered in the structure and represented by a yellow dashed line, The disordered N-terminus is represented with a black dashed line. The other protomer is represented in blue. **(B)** Schematic of ALAS2 chimeras compared to wildtype ALAS2, with C-terminal elements colored as in (A). **(C)** Plot of *in vitro* degradation assays of ALAS2 and C-terminal variants as in Figure 1B with POLDIP2 and hemin (n = 3). **(D)** Representative images of Sypro Red-stained co-IPs of full-length ALAS2 (A2) and C-terminally truncated ALAS2 (A2^ΔC^). Samples were visualized *in* Sypro Red-stained SDS-PAGE gels. (n = 3). **(E)** Degradation of N- or C-terminal SUMO fusions of ALAS2 as in figure 1B with POLDIP2 and hemin (n = 3). Data points with less than 50% depletion of ALAS2 variant were used for the linear fit. The rate of degradation of wt ALAS2 from Fig. 1A is depicted with a black dashed line. See Fig. S5 for representative images of SDS-PAGE gels of degradation time courses.

## Discussion

Here, we outline a mechanism for conditional control of mitochondrial protein degradation. POLDIP2 acts as an adaptor protein that commits ALAS to degradation by CLPXP by a direct heme-induced interaction, thus providing the missing link between CLPX and heme-induced degradation of ALAS. POLDIP2 is crucial both in vitro and in cells for heme-induced degradation of ALAS. By structural modeling and mutagenesis, we identify the N-terminal heme-binding CP motif of ALAS as the site for interaction with POLDIP2. The constitutive association of POLDIP2 with CLPX and its conservation in heme auxotrophs suggests that it controls substrate selection by mitochondrial CLPX more broadly, to be delineated in future studies.

Our spectroscopic observations of heme with ALAS and POLDIP2 indicates that heme binds the ALAS-POLDIP2 complex in a distinctive coordination complex. Mutation of key residues with iron-coordination potential on either side of the complex blocks the observed spectral shift and physical complex formation. These data together are strongly indicative of ALAS-POLDIP2 co-coordination of heme iron, positioning heme to induce ALAS-POLDIP2 interaction in part as a physical component of the protein interface. Further structural characterization of this complex will illuminate how heme acts in this unusual role as a natural molecular glue.

Our analysis of how different elements in ALAS contribute to degradation suggest that in tethering ALAS to CLPX, POLDIP2 may also set the directionality of unfolding. Our data demonstrate that both the N- and C-terminal extensions from the ALAS enzyme core are crucial for heme-induced degradation, but only the N-terminal contact contributes to heme-induced complex assembly, whereas the C-terminus is critical for degradation. The most straightforward model that accounts for these findings is that POLDIP2 first recognizes ALAS2 through a heme-occupied CP motif within its N-terminal element. CLPX then engages ALAS2 within its C-terminal element and initiates unfolding there (coupled to degradation by CLPP). This model also provides a mechanism for how unfolding by CLPX could be diverted from partial unfolding leading to activation (*6*) to complete degradation. CLPX engages ALAS for activation from the N-terminus; by binding the N-terminal extension, POLDIP2 may physically occlude this site, or specifically orient ALAS such that CLPX engages ALAS from the C-terminus. Because the mechanical stability of proteins is directional, unfolding from the C-terminus could allow CLPX to bypass the block that it encounters during activation.

Our findings provide insight into mechanisms of disease and potential targets for treatment in several disorders of heme biosynthesis. Two forms of erythropoietic protoporphyria have been identified that result from misregulation of ALAS2. One is linked to mutations in CLPX that disrupt its degradation of ALAS2 (*3*). Another form is linked to C-terminal truncation or mutation of ALAS2 (XLDPP) (*30*).

XLDPP-linked mutations in ALAS2 that truncate the C-terminal extension not only hyperactivate the enzyme but would also block the mechanism for feedback degradation we delineate here, thus compounding the oversupply of ALA and downstream porphyrin accumulation. Our data suggest that mutations in POLDIP2 should be examined as a possible cause of porphyrias and that therapeutic modulation of this mechanism could be useful in treating multiple porphyrias and anemias.

## MATERIALS AND METHODS

### Cloning of vectors for purification and transient transfection of proteins into cells

Vectors for expression of ALAS2 variants in HEK293 cells were generated with ALAS2 (UniProt P22557, 1-587) with a C-terminal 1D4 epitope tag (*33*) in a pcDNA3.1+ expression vector. Vectors for bacterial expression and purification were generated by cloning the following protein sequences, lacking mitochondrial targeting sequences, and their related variants into pET28b or pET23a with an N-terminal His_6_-SUMO tag: human mitochondrial CLPX (O76031, 65-633), ALAS2 (P22557, 54-587), and POLDIP2 (Q9Y2S7, 52-368). A construct for expressing mitochondrial CLPP (Q16740, 57-277) was previously described (*34*). Standard molecular biology procedures were used.

### Expression and purification of proteins

Constructs were co-transformed into BL21(DE3) *E. coli* for expression in addition to a pRIL plasmid to supplement rare codons. Cultures were grown in Lennox LB at 30°C with shaking at 220 rpm until the desired OD_600_ was achieved (mtCLPX: 0.5, mtCLPP: 0.5, ALAS2: 0.5, POLDIP2: 0.8) at which point cultures were cooled/warmed to the desired expression condition (mtCLPX: 16°C, mtCLPP: 37°C, ALAS2: 16°C, POLDIP2: 18°C), induced with IPTG (mtCLPX: 1 mM, mtCLPP: 0.5 mM, ALAS2: 1 mM, POLDIP2: 0.4 mM), and expressed (mtCLPX, ALAS2, and POLDIP2: overnight; mtCLPP: 3 hours). After expression, cells were harvested by centrifugation at 5000 x g for 10 minutes, washed once in lysis buffer by centrifugation, snap-frozen in liquid nitrogen, and stored at -80°C until purification.

Cell pellets containing ALAS2, mtCLPX, mtCLPP, and POLDIP2 or variants were lysed by two passes through an ice-cold microfluidizer chamber (LM20-30, Microfluidics) in the same base buffer (25 mM HEPES pH 8, 2 mM MgCl_2_, 20 mM Imidazole, 10% glycerol) with varying pressure and salt concentrations (mtCLPX: 10 kpsi, 100 mM KCl and 400 mM NaCl; mtCLPP, ALAS2: 18 kpsi, 100 mM KCl and 400 mM NaCl; POLDIP2: 18 kpsi, 500 mM KCl). All buffers were supplemented prior to use with 1 mM DTT and 0.5 mM PMSF. Buffers for ALAS2 were additionally supplemented with 20 μM PLP. Proteins were captured with HisPur Ni-NTA resin (Thermo Scientific), washed with additional lysis buffer, and then eluted with lysis buffer supplemented to 250 mM Imidizole. Eluted proteins were pooled and incubated with enhanced SUMO Protease (Ulp1_R3, Lau…Bahl JBC 2018) to digest 6H-SUMO tags and dialyzed overnight into their respective storage buffers. All proteins were prepared in the same base storage buffer (25 mM HEPES pH 7.6, 10% Glycerol, 1 mM DTT, 2 mM MgCl_2_) with additional salt (CLPX: 300 mM KCl, CLPP: 100 mM KCl, ALAS2: 150 mM KCl, POLDIP2: 500 mM KCl). Cleaved SUMO tags were removed by Ni-NTA capture. mtCLPX was additionally purified by flowing through Sepharose Q resin. All proteins were purified by gel filtration (mtCLPX, mtCLPP, ALAS2: Superdex S200, POLDIP2: Superdex S75). Fractions corresponding to desired proteins were concentrated, aliquoted, and snap frozen in liquid nitrogen.

### UV-vis spectroscopy

UV-vis spectra were captured from 200-700nm in 5nm steps on a Spectramax 5000 Plate Reader in cuvette mode. Samples contained a final concentration of 25 mM HEPES pH 7.6, 5 mM MgCl_2_, 10% glycerol, 150 mM KCl and 25 μM of each protein and hemin. Hemin was prepared at ≥10 mM in DMSO and diluted in buffer shortly prior to observation. The baseline for each spectrum was set as 0 at 700 nm. The spectrum of each enzyme alone was subtracted from the spectrum of the enzyme incubated with equimolar heme and normalized to absorbance at 270 nm.

### Coprecipitation of purified proteins

Purified proteins were incubated together with anti-FLAG magnetic agarose Beads (Pierce A36797 or Sigma M8823) in 25 mM HEPES pH 7.6, 150 mM KCl, 5 mM MgCl_2_, 10% glycerol, 0.5 mg/mL acetylated BSA, 0.1% Triton X-100, and 1 mM ATPγS. After 45 minute rotating incubation, beads were washed three times in same buffer without acBSA and 0.1 mM ATPγS. For experiments in which CLPX was not included as a variable, ATPγS was omitted. Samples from immunoprecipitation experiments to probe tripartite complex formation with a CLPX-3xFLAG probe were eluted by boiling in Laemmli sample buffer. Samples from immunoprecipitation to directly test POLDIP2 interactions with a POLDIP2-3xFLAG probe were eluted with 0.1 M glycine pH 2.0 for 5 minutes, then neutralized with 1M Tris pH 8.0. All elution samples were separated by SDS-PAGE, stained with SYPRO Red Protein Gel Stain (Invitrogen S6653) at a 1:5000 dilution in 7.5% acetic acid, and imaged on a Typhoon RGB.

### Protein degradation by CLPXP *in vitro*

Degradation of ALAS2 and its variants by CLPXP was monitored *in vitro* by incubating 10 uM ALAS2 with 0.5 µM CLPX (hexamer concentration) and 0.5 µM CLPP in 25 mM HEPES pH 7.6, 150 mM KCl, 5 mM MgCl_2_, 10% glycerol, 1 mM DTT with 4 mM ATP and an ATP regenerating system (5 mM creatine phosphate and 50 mg/mL creatine kinase) at 37°C. 3 µM POLDIP2 and/or 10 µM hemin (Fisher Scientific AAA1116503) were included as indicated. Samples were withdrawn at times indicated and immediately mixed with Laemmli SDS sample buffer. Samples were heat-denatured, separated by SDS-PAGE, and stained with SYPRO Red. Gels were imaged on an Amersham Typhoon RGB scanner and quantified in ImageQuant using rolling-ball background subtraction. Rates were extracted using linear fits to all replicates in GraphPad Prism.

### Construction of HEK293 POLDIP2 knockout line

Synthetic complementary oligos targeting POLDIP2 (5’-CACCGCGCGTCGTCGTGGTCGACGC-3’ and 5’-AAACGCGTCGACCACGACGACGCGC -3’) were cloned into PX459-SpCas9 (Addgene, (*35*)) at the BbsI site. The resulting plasmid was transfected into HEK293 cells at 70% confluency using Genejuice (Fisher Scientific 70-967-3) at a 1:3 ratio (DNA : Genejuice). 24 hours post-transfection, culture media was changed. 48 hours post-transfection, cells were diluted into 96-well plates for a final approximate cell density of 0.5 cells/well. Wells were monitored and those that contained monoclonal populations were expanded. Knockout was confirmed with Western blotting and Sanger sequencing. Cell lysate was analyzed with an anti-POLDIP2 antibody (Abcam ab68663). Genomic DNA from cell samples was amplified with primers 5’-GGTCGGGCTCTGTGTCAG -3’ 5’-AGGGCTGAATTAAAAAGCTTCC-3’ to analyze the cut site with Sanger sequencing.

### Monitoring of ALAS degradation in cells

HEK293 cells (CRL-1573, ATCC) were grown in Dulbecco’s Modified Eagle Medium supplemented with 10% FBS, 1% non-essential amino acids, 4.5 g/L D-glucose, 1 mM sodium pyruvate, and 862.0 mg/L L-alanyl-glutamine (GlutaMAX)) (Gibco). To monitor degradation of ALAS2 variants, cells were transfected 48 hours prior to observation with pcDNA3.1-ALAS2-1D4 or pcDNA3.1-ALAS2(C70S)-1D4 with GeneJuice transfection reagent (Sigma-Aldrich). To monitor degradation of endogenous ALAS1, cells were plated 72 hours prior to observation and succinylacetone (MedChemExpress HY-W010184) was added to 0.5 mM 24 hours prior to observation. To initiate the observation, 100 µg/mL cycloheximide and 100 µM hemin (in DMSO) or DMSO alone were added to the cells. 0-hour timepoints were not supplied with cycloheximide, hemin, or DMSO. Total cell volume was collected from one well per time point, and all subsequent collection steps were performed at 4°C. Cell pellets were washed with PBS and lysed in RIPA lysis buffer (50 mM Tris base pH 8, 150 mM NaCl, 0.1% SDS, 0.5% sodium deoxycholate, 1% Triton X 100, and 0.5 mM PMSF) before analysis by Western blot.

### Western blotting

Samples was separated by SDS-PAGE and transferred to PVDF. Membranes were blocked with 1.5% bovine serum albumin (BSA) in TBS with 0.05% Tween 20 (TBST) and probed with primary antibodies, source and working dilution as follows: anti-1D4 (made in-house) 0.2 µg/mL; anti-ALAS1, Abcam ab154860, 1:5000; anti-β-actin antibody, Santa Cruz Biotechnology sc-47778, 1:1000. Membranes were subsequently probed with fluorescently-conjugated secondary antibodies: anti-mouse Dylight 680-conjugated (Thermo Scientific 35519,1:5000 dilution) or anti-rabbit IRDye 800CW-conjugated (LI-COR 92632211, 1:10000 dilution).

## ACKNOWLEDGEMENTS

We would like to acknowledge N. Bradshaw and M. Fahie for valuable discussions and critical feedback on the manuscript. Funding was provided by National Institutes of Health grant R01GM151332 (J.R.K), T32GM139798 (T.C.) and T32GM007596 (C.P. and A.D.).

---

**Supplementary Figure 1:**
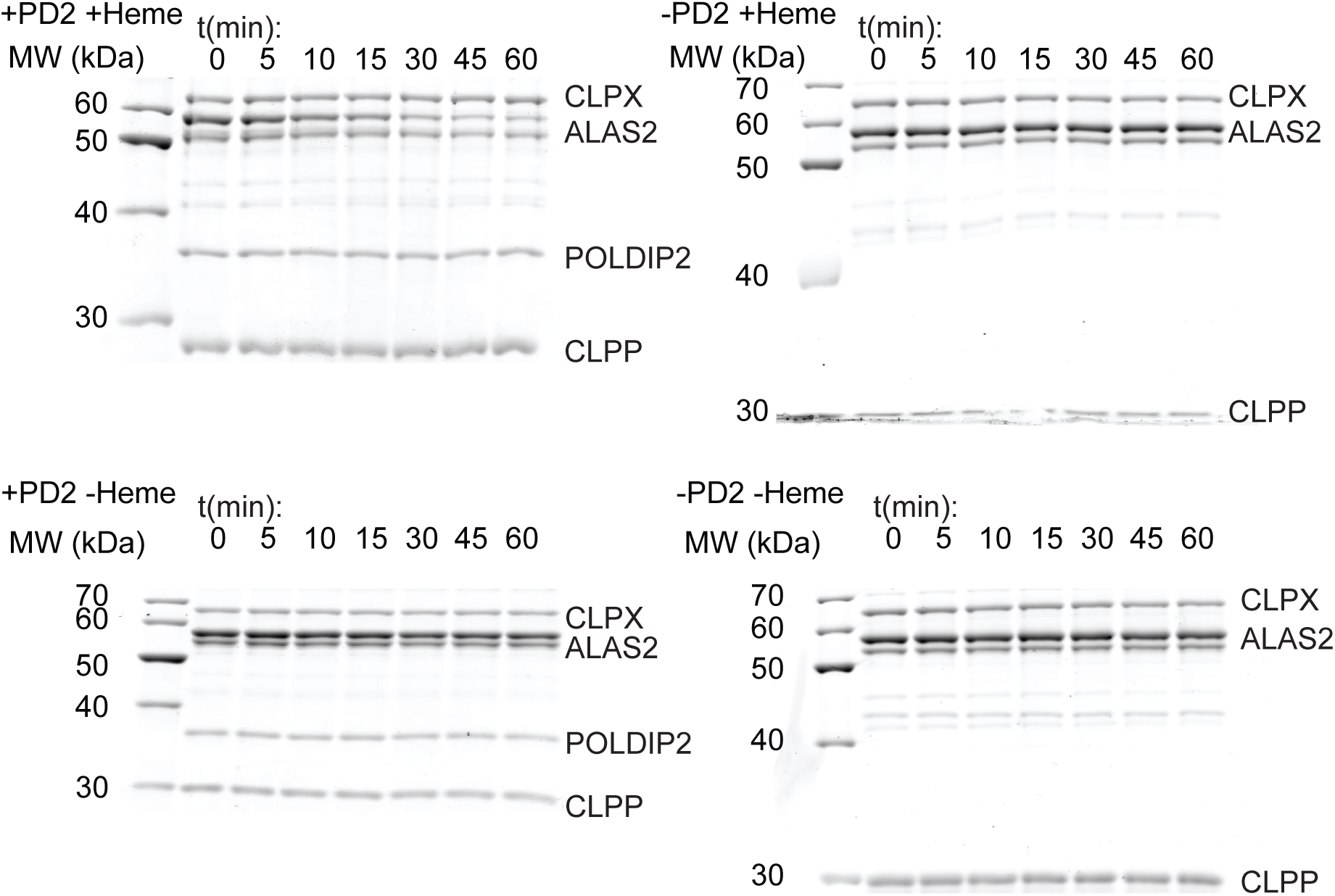
Representative images of ALAS2 degradation by CLPXP. 10 µM ALAS2 was incubated with 0.5 µM CLPX and CLPP (hexamer and 14-mer respectively), 4 mM ATP, +/-3 µM POLDIP2, +/-10 µM hemin (H). (n = 5 for +hemin +PD2, 3 for other conditions).

**Supplementary Figure 2:**
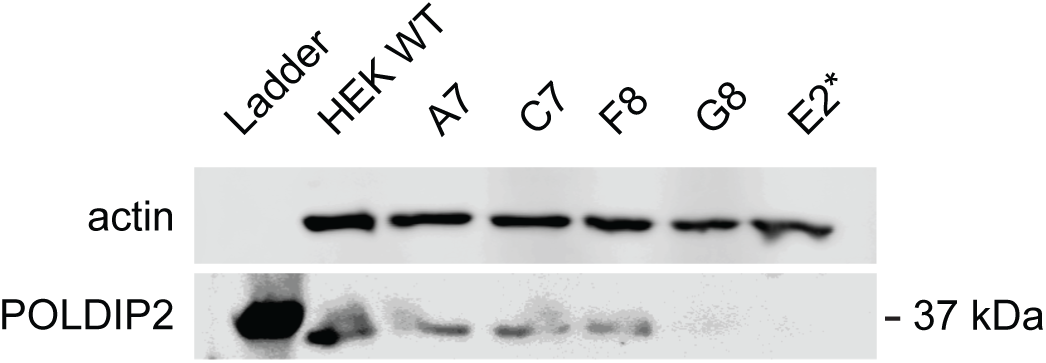
Validation of a POLDIP2-knockout HEK293 cell line. Western blot to screen HEK293 monoclonal populations after transfection with PX459-SpCas9 vector with gRNA targeting the POLDIP2 gene. 5 wells were screened (A7, C7, F8, G8, and E2) and compared with wildtype cells. E2 was chosen as the final knockout line.

**Supplementary Figure 3:**
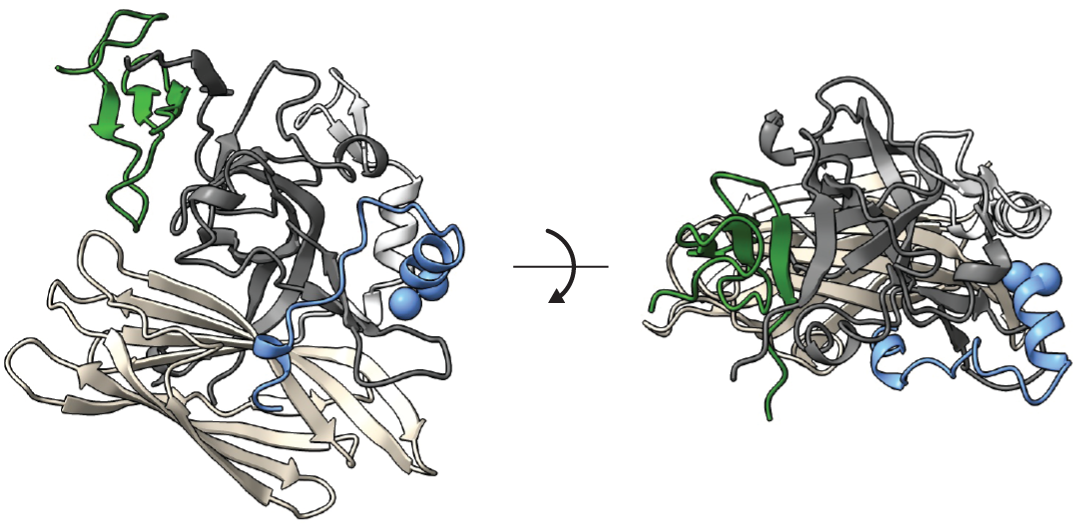
AlphaFold2-multimer model of an ALAS2-POLDIP2-CLPX complex. The interacting zinc-finger N-domain of CLPX is shown in green. The interacting peptide of ALAS2 is shown in blue, with CP70-P71 highlighted with blue spheres at the α carbon. POLDIP2 is shown in dark gray (YccV domain), light gray (linker), and wheat (ApaG domain).

**Supplementary Figure 4:**
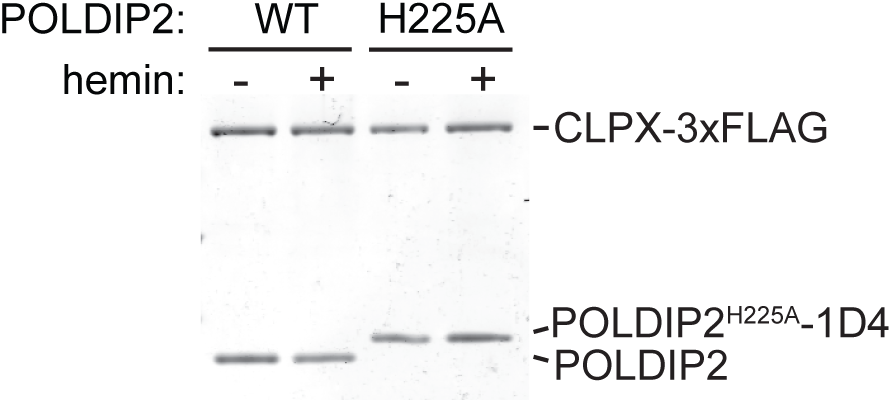
H225A mutation in POLDIP2 does not perturb interaction with CLPX. Coimmunoprecipitation of 5 µM POLDIP2 (wt or H225A) with 0.5 µM CLPX-3xFLAG (hexamer) +/-10 µM hemin. Samples were visualized by Sypro Red-stained SDS-PAGE gels. Representative image shown; n = 2.

**Supplementary Figure 5:**
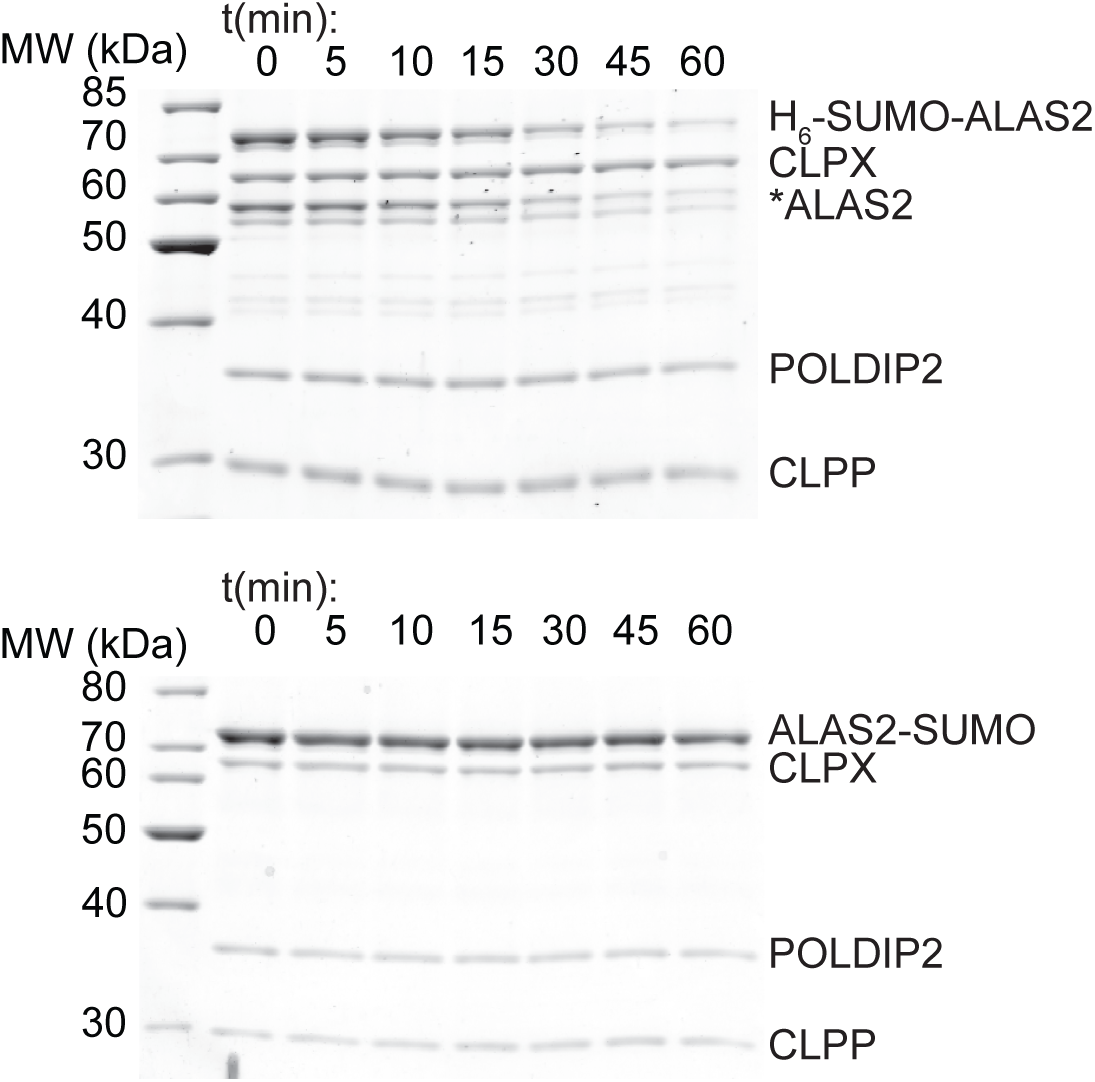
Representative images of degradation time courses of N- and C-terminal SUMO fusions with ALAS2. 10 µM of 6H-SUMO-ALAS2 or ALAS2-SUMO were incubated with 0.5 µM CLPX and CLPP (hexamer and 14-mer respectively), 4mM ATP, 3 µM POLDIP2, and 10 µM hemin. H_6_SUMO-ALAS2 copurified with a truncation product, approximately the size of untagged ALAS2. Only the full-length protein was quantified for displayed in Fig. 4. N = 3 for each chimera.

## REFERENCES

1. E. M. Jenkinson, A. U. Rehman, T. Walsh, J. Clayton-Smith, K. Lee, R. J. Morell, M. C. Drummond, S. N. Khan, M. A. Naeem, B. Rauf, N. Billington, J. M. Schultz, J. E. Urquhart, M. K. Lee, A. Berry, N. A. Hanley, S. Mehta, D. Cilliers, P. E. Clayton, H. Kingston, M. J. Smith, T. T. Warner, U. of W. C. for M. Genomics, G. C. Black, D. Trump, J. R. E. Davis, W. Ahmad, S. M. Leal, S. Riazuddin, M.-C. King, T. B. Friedman, W. G. Newman, Perrault syndrome is caused by recessive mutations in CLPP, encoding a mitochondrial ATP-dependent chambered protease. Am J Hum Genet 92, 605–613 (2013).

2. A. Cole, Z. Wang, E. Coyaud, V. Voisin, M. Gronda, Y. Jitkova, R. Mattson, R. Hurren, S. Babovic, N. Maclean, I. Restall, X. Wang, D. V. Jeyaraju, M. A. Sukhai, S. Prabha, S. Bashir, A. Ramakrishnan, E. Leung, Y. H. Qia, N. Zhang, K. R. Combes, T. Ketela, F. Lin, W. A. Houry, A. Aman, R. Al-awar, W. Zheng, E. Wienholds, C. J. Xu, J. Dick, J. C. Y. Wang, J. Moffat, M. D. Minden, C. J. Eaves, G. D. Bader, Z. Hao, S. M. Kornblau, B. Raught, A. D. Schimmer, Inhibition of the Mitochondrial Protease ClpP as a Therapeutic Strategy for Human Acute Myeloid Leukemia. Cancer Cell 27, 864–876 (2015).

3. Y. Y. Yien, S. Ducamp, L. N. van der Vorm, J. R. Kardon, H. Manceau, C. Kannengiesser, H. A. Bergonia, M. D. Kafina, Z. Karim, L. Gouya, T. A. Baker, H. Puy, J. D. Phillips, G. Nicolas, B. H. Paw, Mutation in human CLPX elevates levels of δ-aminolevulinate synthase and protoporphyrin IX to promote erythropoietic protoporphyria. Proceedings of the National Academy of Sciences 114, E8045– E8052 (2017).

4. A. Belot, H. Puy, I. Hamza, H. L. Bonkovsky, Update on heme biosynthesis, tissue-specific regulation, heme transport, relation to iron metabolism and cellular energy. Liver Int., doi: 10.1111/liv.15965 (2024).

5. K. Peoc’h, G. Nicolas, C. Schmitt, A. Mirmiran, R. Daher, T. Lefebvre, L. Gouya, Z. Karim, H. Puy, Regulation and tissue-specific expression of δ-aminolevulinic acid synthases in non-syndromic sideroblastic anemias and porphyrias. Mol Genet Metab 128, 190–197 (2019).

6. J. R. Kardon, J. A. Moroco, J. R. Engen, T. A. Baker, Mitochondrial ClpX activates an essential biosynthetic enzyme through partial unfolding. eLife 9 (2020).

7. A. J. R. Kardon, Y. Y. Yien, N. C. Huston, D. S. Branco, G. J. Hildick-Smith, K. Y. Rhee, B. H. Paw, T. A. Baker, Mitochondrial ClpX Activates a Key Enzyme for Heme Biosynthesis and Erythropoiesis. Cell 161, 858–867 (2015).

7. Y. Kubota, K. Nomura, Y. Katoh, R. Yamashita, K. Kaneko, K. Furuyama, Novel Mechanisms for Heme-dependent Degradation of ALAS1 Protein as a Component of Negative Feedback Regulation of Heme Biosynthesis. Journal of Biological Chemistry, doi: 10.1074/jbc.m116.719161 (2016).

8. N. J. Kuhlmann, P. Chien, Selective adaptor dependent protein degradation in bacteria. Curr. Opin. Microbiol. 36, 118–127 (2017).

9. A. A. Kulik, K. K. Maruszczak, D. C. Thomas, N. L. A. Nabi-Aldridge, M. Carr, R. J. Bingham, C. D. O. Cooper, Crystal structure and molecular dynamics of human POLDIP2, a multifaceted adaptor protein in metabolism and genome stability. Protein Sci. 30, 1196–1209 (2021).

10. F. Paredes, K. Sheldon, B. Lassègue, H. C. Williams, E. A. Faidley, G. A. Benavides, G. Torres, F. Sanhueza-Olivares, S. M. Yeligar, K. K. Griendling, V. Darley-Usmar, A. S. Martin, Poldip2 is an oxygen-sensitive protein that controls PDH and αKGDH lipoylation and activation to support metabolic adaptation in hypoxia and cancer. Proceedings of the National Academy of Sciences 115, 1789–1794 (2018).

11. P. R. Strack, E. J. Brodie, H. Zhan, V. J. Schuenemann, L. J. Valente, T. Saiyed, B. R. Lowth, L. M. Angley, M. A. Perugini, K. Zeth, K. N. Truscott, D. A. Dougan, Polymerase delta-interacting protein 38 (PDIP38) modulates the stability and activity of the mitochondrial AAA+ protease CLPXP. Communications Biology 3, 646–12 (2020).

12. K. Luck, D.-K. Kim, L. Lambourne, K. Spirohn, B. E. Begg, W. Bian, R. Brignall, T. Cafarelli, F. J. Campos-Laborie, B. Charloteaux, D. Choi, A. G. Coté, M. Daley, S. Deimling, A. Desbuleux, A. Dricot, M. Gebbia, M. F. Hardy, N. Kishore, J. J. Knapp, I. A. Kovács, I. Lemmens, M. W. Mee, J. C. Mellor, C. Pollis, C. Pons, A. D. Richardson, S. Schlabach, B. Teeking, A. Yadav, M. Babor, D. Balcha, O. Basha, C. Bowman-Colin, S.-F. Chin, S. G. Choi, C. Colabella, G. Coppin, C. D’Amata, D. D. Ridder, S. D. Rouck, M. Duran-Frigola, H. Ennajdaoui, F. Goebels, L. Goehring, A. Gopal, G. Haddad, E. Hatchi, M. Helmy, Y. Jacob, Y. Kassa, S. Landini, R. Li, N. van Lieshout, A. MacWilliams, D. Markey, J. N. Paulson, S. Rangarajan, J. Rasla, A. Rayhan, T. Rolland, A. San-Miguel, Y. Shen, D. Sheykhkarimli, G. M. Sheynkman, E. Simonovsky, M. Taşan, A. Tejeda, V. Tropepe, J.-C. Twizere, Y. Wang, R. J. Weatheritt, J. Weile, Y. Xia, X. Yang, E. Yeger-Lotem, Q. Zhong, P. Aloy, G. D. Bader, J. D. L. Rivas, S. Gaudet, T. Hao, J. Rak, J. Tavernier, D. E. Hill, M. Vidal, F. P. Roth, M. A. Calderwood, A reference map of the human binary protein interactome. Nature 580, 402–408 (2020).

13. H. J. Bailey, G. A. Bezerra, J. R. Marcero, S. Padhi, W. R. Foster, E. Rembeza, A. Roy, D. F. Bishop, R. J. Desnick, G. Bulusu, H. A. Dailey, W. W. Yue, Human aminolevulinate synthase structure reveals a eukaryotic-specific autoinhibitory loop regulating substrate binding and product release. Nat Comms 11, 1–12 (2020).

14. D. F. Bishop, V. Tchaikovskii, I. Nazarenko, R. J. Desnick, Molecular expression and characterization of erythroid-specific 5-aminolevulinate synthase gain-of-function mutations causing X-linked protoporphyria. Mol. Med. 19, 18–25 (2013).

15. Q. Tian, T. Li, W. Hou, J. Zheng, L. Schrum, H. Bonkovsky, LONP1-dependent breakdown of mitochondrial 5-aminolevulinic acid synthase protein by heme in human liver cells. Journal of Biological Chemistry 286, 26424–26430 (2011).

16. K. Nomura, Y. Kitagawa, M. Aihara, Y. Ohki, K. Furuyama, T. Hirokawa, Heme-dependent recognition of 5-aminolevulinate synthase by the human mitochondrial molecular chaperone ClpX. Febs Lett 595, 3019–3029 (2021).

17. M. Mirdita, K. Schütze, Y. Moriwaki, L. Heo, S. Ovchinnikov, M. Steinegger, ColabFold: making protein folding accessible to all. Nat Methods 19, 679–682 (2022).

18. J. Yeom, E. A. Groisman, Activator of one protease transforms into inhibitor of another in response to nutritional signals. Genes Dev 33, 1280–1292 (2019).

19. N. Puri, A. W. Karzai, HspQ Functions as a Unique Specificity-Enhancing Factor for the AAA+ Lon Protease. Molecular Cell 66, 672–683.e4 (2017).

20. R. K. Mallampalli, T. A. Coon, J. R. Glasser, C. Wang, S. R. Dunn, N. M. Weathington, J. Zhao, C. Zou, Y. Zhao, B. B. Chen, Targeting F Box Protein Fbxo3 To Control Cytokine-Driven Inflammation. J. Immunol. 191, 5247–5255 (2013).

21. Z. Yu, T. Chen, X. Li, M. Yang, S. Tang, X. Zhu, Y. Gu, X. Su, M. Xia, W. Li, X. Zhang, Q. Wang, X. Cao, J. Wang, Lys29-linkage of ASK1 by Skp1−Cullin 1−Fbxo21 ubiquitin ligase complex is required for antiviral innate response. eLife 5, e14087 (2016).

22. K. Nishimura, J. Apitz, G. Friso, J. Kim, L. Ponnala, B. Grimm, K. J. van Wijk, Discovery of a Unique Clp Component, ClpF, in Chloroplasts: A Proposed Binary ClpF-ClpS1 Adaptor Complex Functions in Substrate Recognition and Delivery. THE PLANT CELL 27, 2677–2691 (2015).

23. A. S. Richter, C. Banse, B. Grimm, The GluTR-binding protein is the heme-binding factor for feedback control of glutamyl-tRNA reductase. eLife 8 (2019).

24. H. Cheng, R. D. Schaeffer, Y. Liao, L. N. Kinch, J. Pei, S. Shi, B.-H. Kim, N. V. Grishin, ECOD: An Evolutionary Classification of Protein Domains. PLoS Comput. Biol. 10, e1003926 (2014).

25. E. Y. Park, B.-G. Lee, S.-B. Hong, H.-W. Kim, H. Jeon, H. K. Song, Structural Basis of SspB-tail Recognition by the Zinc Binding Domain of ClpX. J. Mol. Biol. 367, 514–526 (2007).

26. H. H. Brewitz, T. Kühl, N. Goradia, K. Galler, J. Popp, U. Neugebauer, O. Ohlenschläger, D. Imhof, Role of the Chemical Environment beyond the Coordination Site: Structural Insight into FeIII Protoporphyrin Binding to Cysteine-Based Heme-Regulatory Protein Motifs. ChemBioChem 16, 2216– 2224 (2015).

27. R. T. Sauer, T. A. Baker, AAA+ proteases: ATP-fueled machines of protein destruction. Annu Rev Biochem 80, 587–612 (2011).

28. S. D. Whatley, S. Ducamp, L. Gouya, B. Grandchamp, C. Beaumont, M. N. Badminton, G. H. Elder, S. A. Holme, A. V. Anstey, M. Parker, A. V. Corrigall, P. N. Meissner, R. J. Hift, J. T. Marsden, Y. Ma, G. Mieli-Vergani, J.-C. Deybach, H. Puy, C-terminal deletions in the ALAS2 gene lead to gain of function and cause X-linked dominant protoporphyria without anemia or iron overload. Am J Hum Genet 83, 408–414 (2008).

29. S. Ducamp, X. Schneider-Yin, F. de Rooij, J. Clayton, E. J. Fratz, A. Rudd, G. Ostapowicz, G. Varigos, T. Lefebvre, J.-C. Deybach, L. Gouya, P. Wilson, G. C. Ferreira, E. I. Minder, H. Puy, Molecular and Functional Analysis of the C-Terminal Region of Human Erythroid Specific 5-Aminolevulinic Synthase Associated with X-Linked Dominant Protoporphyria (XLDPP). Hum Mol Genet, doi: 10.1093/hmg/dds531 (2012).

30. G. A. Hunter, G. C. Ferreira, An Extended C-Terminus, the Possible Culprit for Differential Regulation of 5-Aminolevulinate Synthase Isoforms. Frontiers Mol Biosci 9, 920668 (2022).

31. S. Kadirvel, K. Furuyama, H. Harigae, K. Kaneko, Y. Tamai, Y. Ishida, S. Shibahara, The carboxyl-terminal region of erythroid-specific 5-aminolevulinate synthase acts as an intrinsic modifier for its catalytic activity and protein stability. Exp Hematol 40, 477–486.e1 (2012).

32. D. MacKenzie, A. Arendt, P. Hargrave, J. H. McDowell, R. S. Molday, Localization of binding sites for carboxyl terminal specific anti-rhodopsin monoclonal antibodies using synthetic peptides. Biochemistry 23, 6544–6549 (1984).

33. S. G. Kang, J. Ortega, S. K. Singh, N. Wang, N.-N. Huang, A. C. Steven, M. R. Maurizi, Functional proteolytic complexes of the human mitochondrial ATP-dependent protease, hClpXP. J Biol Chem 277, 21095–21102 (2002).

34. F. A. Ran, P. D. Hsu, J. Wright, V. Agarwala, D. A. Scott, F. Zhang, Genome engineering using the CRISPR-Cas9 system. Nature Protocols 8, 2281–2308 (2013).

